# Impact of rare structural variant events in newly diagnosed multiple myeloma

**DOI:** 10.1101/2023.01.03.522573

**Authors:** Monika Chojnacka, Benjamin Diamond, Bachisio Ziccheddu, Even Rustad, Kylee Maclachlan, Marios Papadimitriou, Eileen M. Boyle, Patrick Blaney, Saad Usmani, Gareth Morgan, Ola Landgren, Francesco Maura

## Abstract

Whole genome sequencing (WGS) of newly diagnosed multiple myeloma patients (NDMM) has shown recurrent structural variant (SV) involvement in distinct regions of the genome (i.e. hotspots) and causing recurrent copy number alterations. Together with canonical immunoglobulin translocations, these SVs are recognized as “recurrent SVs”. More than half SVs were not involved in recurrent events. The significance of these “rare SVs” has not been previously examined. In this study, we utilize 752 WGS and 591 RNA-seq data from NDMM patients to determine the role of rare SVs in myeloma pathogenesis. 94% of patients harbored at least one rare SV event. Rare SVs showed an SV-class specific enrichment within genes and superenhancers associated with outlier gene expression. Furthermore, known myeloma driver genes recurrently impacted by point mutations were dysregulated by rare SVs. Overall, we demonstrate the association of rare SVs with aberrant gene expression supporting a driver role in myeloma pathogenesis.

**SIGNIFICANCE:** Characterization of multiple myeloma genome revealed that more than half structural variants are not involved in recurrent events. Here, we demonstrate that these rare SVs hold potential for myeloma pathogenesis through their gene expression impact. Rare SVs contribute to MM heterogeneity and have implications for development of individualized treatment.

## INTRODUCTION

Whole genome sequencing (WGS) has shown that the clinical behavior of various cancer types is driven by a complex genomic landscape characterized by multiple genomic drivers and mutational processes[1, 2]. Among these somatic events, structural variants (SV) have emerged as key events causing the acquisition of copy number alterations (CNA), creating gene fusions, and dysregulating gene expression through superenhancer hijacking and the disruption of 3D genomic structure[1, 3-6]. A single SV is comprised of 2 breakpoints and classified according to four main classes: duplication, deletion, inversion, and translocation. Each SV unit may exist as a singleton or as part of a complex SV event – such as chromothripsis, templated insertion, or chromoplexy, and occur at various rates across cancer types[1, 3, 7].

Multiple myeloma (MM) is a clinically and biologically heterogenous hematological malignancy characterized by an expansion of abnormal plasma cells in the bone marrow[6, 8]. Historically, SV analysis in MM has been restricted to recurrent translocations between immunoglobulin genes (Ig) and key oncogenes such as *CCND1, MAF* and *NSD2*. These “Canonical Translocation” events are considered to be one of few initiating genomic myeloma drivers due to their clonal characteristic of conservation across several evolutionary phases of tumor development, their strong impact on gene expression and consequent impact on disease biology [9, 10].

Analysis of large datasets of clinically annotated WGS paired with RNA sequencing (RNA-seq) have allowed the interrogation of SVs within the MM genomic landscape. Multiple studies have shown a heavy SV enrichment in MM beyond the canonical SV events, which account for only a minor fraction of the full SV repertoire [11-14]. Interestingly, the MM SV landscape shows a high prevalence of chromothripsis and chromoplexy, and has one of the highest prevalence of templated insertions described in cancer [12, 15]. More than 60 recurrent SV genomic hotspot have been reported and linked to multiple driver genes and regulatory regions (i.e., “SV Hotspot”)[12]. In addition to these recurrent events, SVs outside of SV hotspots and canonical Ig events are often responsible for recurrent copy number alterations in 152 previously defined genomic regions, here named “GISTIC CNA” after the method of discovery [12]. Despite this comprehensive characterization of SVs to date, more than half SV are not involved in a recurrent “Canonical Translocation”, “SV Hotspot” or “GISTIC CNA” events. It is largely unknown if these unclassified SV events, here defined as “rare SV”, play a driver or passenger role in MM disease biology and whether we can gain further insights into recurrently impacted pathways that could constitute novel targets for therapy. Overall, the relevance of rare SV events is not new in MM. In fact, several rare events have been reported and linked to distinct clinical behavior, like rare nonsynonymous mutations on *CRBN, IKFZ3*, or *BCMA* deletions after exposure to immunomodulary agents and CAR-T, respectively [16, 17]. In this work, to expand our understanding on the role of rare SVs in relation to clinical and biological heterogeneity, and potential implications on the development of individualized treatment strategies, we utilize 752 WGS and 591 RNA-seq data in a newly diagnosed MM (NDMM) patient cohort, demonstrating that many rare SV events are not passengers but rather have a major impact on the gene expression and tumor biology in individual patients.

## RESULTS

### Structural variant and complex event annotation

To characterize the patterns and biological impact of rare SVs in NDMM, we interrogated 752 WGS samples from patients enrolled in the CoMMpass study (NCT01454297; IA13), 591 of which had patient matched RNA-seq data available. To classify an individual SV and complex SV event, we devised a hierarchical ranking system. First, individual SVs were classified based on the highest priority breakpoint involvement. We annotated each individual SV (deletion, duplication, inversion, and translocation) as either rare or recurrent based on the region of the genome the breakpoints occur. Likewise, in the context of complex SV events, the highest priority SV within an event classified the whole SV event. Recurrent SVs included all canonical SV translocations, SVs that fell within SV hotspots, and SVs involved in recurrent GISTIC CNA regions altering copy number, and appropriately classified by the recurrent event or region the SV breakpoint involved (**Fig 1A**; **Methods**) [12]. Any SV with both breakpoints not included in any of these three groups was classified as rare. Hierarchical annotation prioritized canonical translocations, followed by SV hotspots, GISTIC CNA, and finally rare SV events. We focused on 4 main complex events – chromothripsis, templated insertions, chromoplexy and finally bulk categorizing all remaining complex events as “complex”. Reasonably, a complex event annotated as a recurrent SV can be composed of both recurrent and rare SVs (**Fig 1B-E**). In contrast, a fully rare complex SV event will be composed only by rare SVs. Despite main drivers being involved by recurrent SVs, rare SVs can play an additive role, targeting genes with biological impact. In this work we investigated rare SVs in two contexts: 1) rare SVs within recurrent SV events, and 2) rare SVs within fully rare SV events.

**Figure 1:**
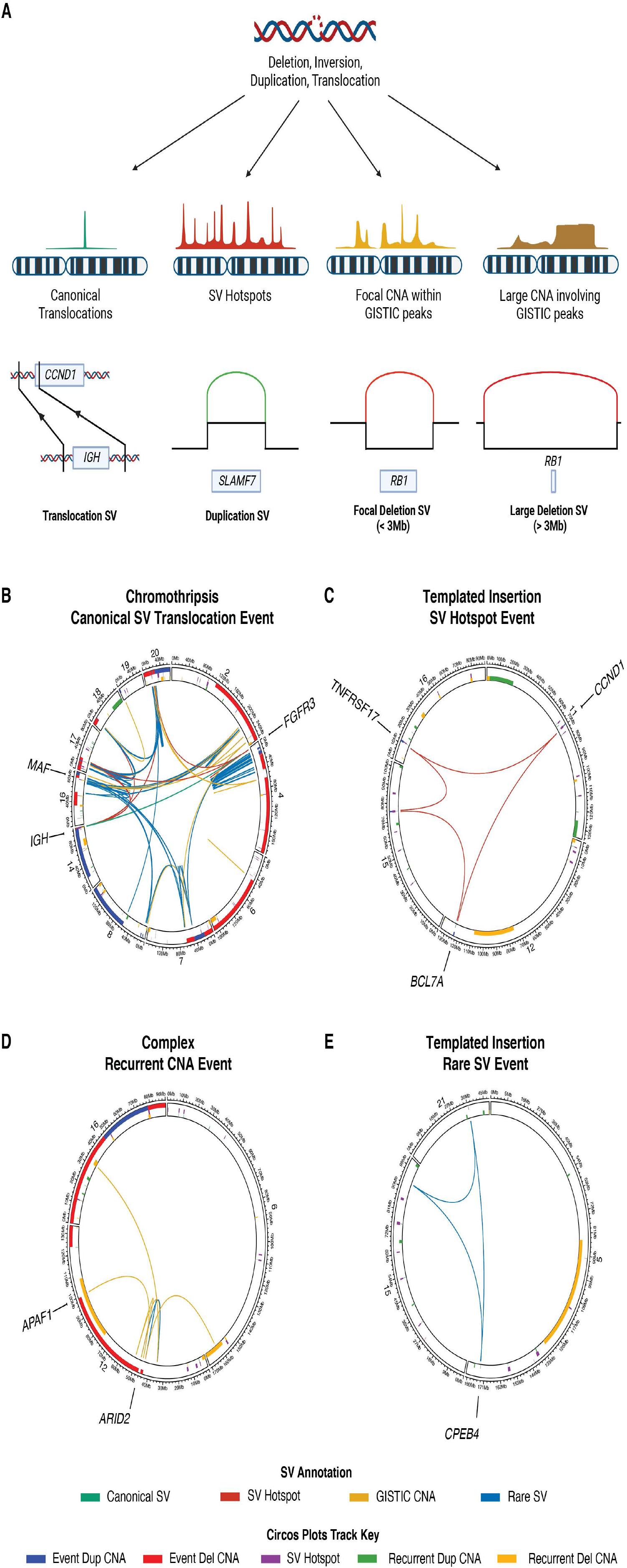
Defining recurrent and rare SV events. **A)** Any SV events which causes a canonical myeloma translocation, or SV breakpoint that falls within the previously defined recurrent regions was identified as a recurrent SV. Examples depict SV breakpoints falling within SV hotspots, and causing focal and large copy umber gains or losses occurring within previously defined recurrent GISTIC CNA regions of the myeloma genome. **B)** Chromothripsis class SV annotated as “Canonical Translocation” event based on the highest priority SV involving a translocation between *IGH* and *FGFR3*/*NSD2* (green) comprising of 123 individual SVs, consisting of 1 canonical translocation between *IGH* and *FGFR3/NSD2*, 17 SVs within hotspot regions, 12 SVs within recurrent GISTIC CNA regions, and 91 rare SVs. **C)** Templated insertion class SV annotated as an “SV Hotspot” event, based on the highest priority SV (red) involving a hotspot region (purple). **D)** Complex SV annotated as a “Recurrent CNA” event based on the highest priority SV involving a recurrent GISTIC deletion CNA region (yellow). **E)** Templated insertion class SV annotated as a “Rare SV” event, based on the exclusion of any recurrent regions affected by the event’s SVs (blue).

### Defining the landscape and role of rare SV and rare SV events in NDMM

At least one rare SV event was observed across 94% (705/752) of all patients (**Fig 2A, Suppl Fig 1A**). Of the total 8,942 SV events across 752 patients, 269 (3%) were classified as Canonical Translocation SVs (128 *CCND1*, 8 *CCND2/CCND3*, 22 *MAF/A/B*, 87 *NSD2*, 24 other Ig translocations), 1,871 (21%) as SV hotspot, and 1,843 (20%) as GISTIC CNA SVs, leaving 4,959 (55%) fully rare SV events (**Fig 2A-B**). Amongst the main MM cytogenetic groups, patients with canonical translocations involving *CCND1* showed significantly lower counts and proportions of rare SVs per event compared to *MAF/A/B, NSD2*, hyperdiploid and other translocation groups (p<0.001, using Wilcoxon test; **Fig 2A, Suppl Fig 1B, Suppl Tables 1-2**). Additionally, patients with *NSD2* translocations had considerably higher counts of rare SV per event compared to all other groups except the *CCND2/CCND3* group (p<0.01, using Wilcoxon test; **Suppl Fig 1B, Suppl Table 1**).

**Figure 2:**
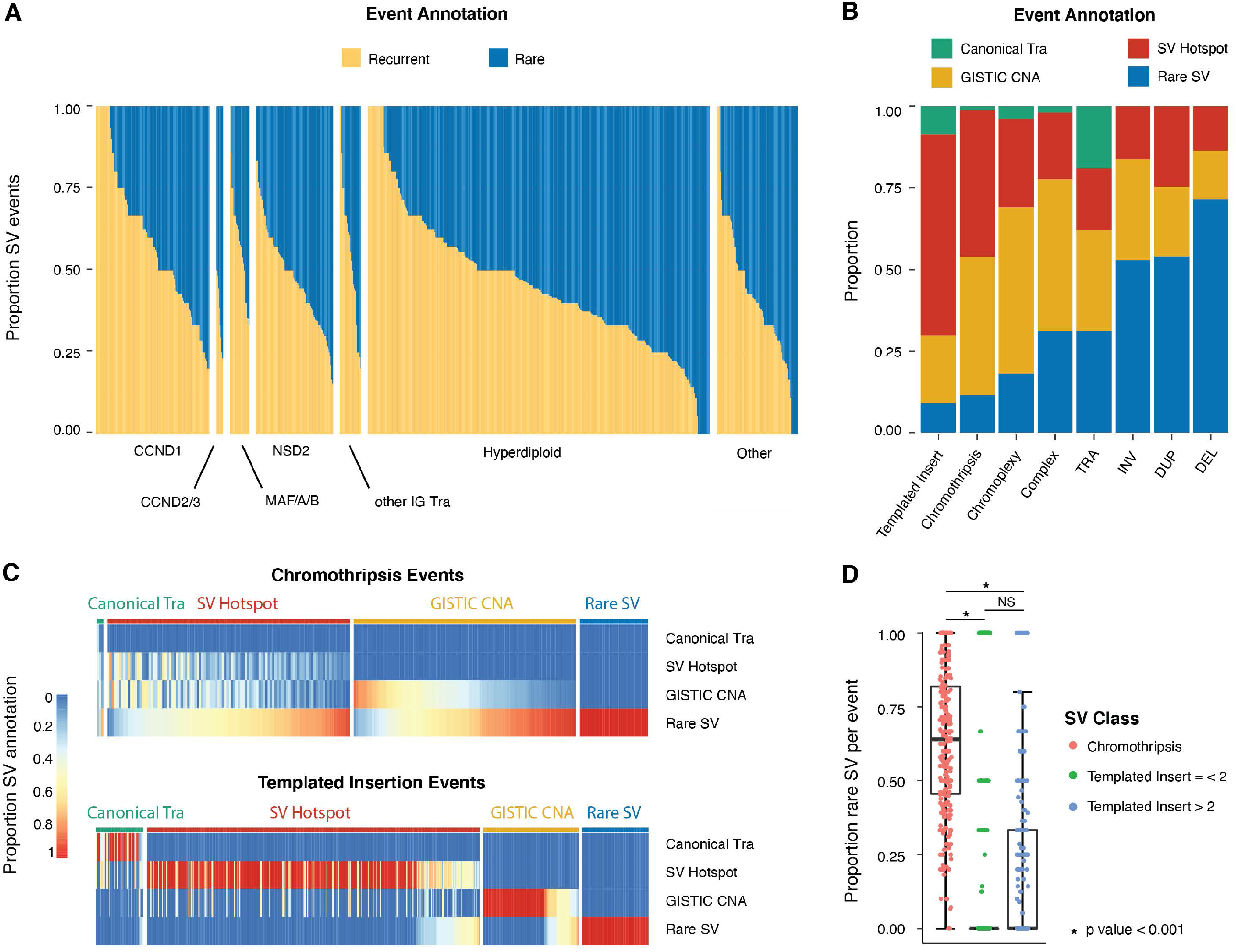
Rare SV landscape of newly diagnosed multiple myeloma patients. **A)** Proportion of rare and recurrent SV events per patient, subdivided into classical clinical cytogenetic groups. *CCND1, CCND2/3, MAF/A/B*, and *NSD2* denote canonical translocations between immunoglobulin (Ig) loci. Group *MAF/A/B* includes genes *MAF, MAFA*, and *MAFB*. “other IG Tra” group represents the cytogenetic group with Ig translocations between *Ig* and non-canonical partners. “Other” cytogenetic group denotes patients without canonical translocations, translocations involving the Ig locus, or a hyperdiploid genome. **B)** Proportion of SV events in each classification category based on annotation hierarchy as described, and grouped by SV class. **C)** Breakdown of SV annotation within each SV event, in chromothripsis and templated insertion complex SV events. **D)** Statistical comparison of proportion of rare Ss per event in all chromothripsis events compared to templated insertion events involving up to 2 chromosomes and, templated insertion involving more than 2 chromosomes (p< 0.001, using Wilcoxon test).

Overall, 42.7% (8,148/19,055) of individual SVs were directly involved in recurrent regions while 57.3% (10,907/19,055) individual SVs occurred in rare unclassified regions. Among rare SVs, 57.3% (6,256/10,907) occurred within fully rare SV events, while 4,651 (42.7%) occurred within recurrent SV events (i.e. complex events composed of both rare and recurrent SV; **Suppl Fig 1C**). Across SV classes, 11% (63/547) of templated insertions, 12% (29/236) of chromothripsis, 18% (18/100) of chromoplexy, and 33% (159/471) of all other complex events were annotated as fully rare SV events. Among the single SV event classes, the ratio between rare and recurrent SV, was lowest for translocations (31%; 334/1,069), followed by inversions (52%; 507/973) and duplications (53%; 886/1,661), and was highest among deletion (71%; 2,782/3,885) – with more rare deletions than in all recurrent class deletions combined (**Fig 2B, Suppl Table 3**).

Considering their impact on the genome and clinical outcome, chromothripsis and templated insertions were of particular interest [12, 18]. A significantly higher proportion of rare SVs were found within chromothripsis events compared to both templated insertions involving two chromosomes and templated insertions involving more than 2 chromosomes, with median rare SV proportions of 64% (0-100% range), 0% (0-100% range), and 0% (0-100% range) of each complex SV event, respectively (p<0.001, using Wilcoxon test; **Fig 2C-D, Suppl Fig 1D–E**). The differences observed between chromothripsis and templated insertions is expected, considering that the latter tends to be associated with focal amplification of recurrent driver genes and regulatory regions, while chromothripsis is often composed of multiple SVs affecting broad genomic segments often across multiple chromosomes. Chromoplexy and other complex events also displayed a large proportion of rare SVs within recurrent events, with median rare SV proportion of 55% (16-100% range) and 75% (12-100% range), each significantly higher than templated insertions (p<0.001; using Wilcoxon test; **Suppl Fig 1D-E**). Having established that rare SVs occur in most of the patients and across all SV classes, we first focused on rare SVs targeting myeloma genes of known impact.

### Rare SV as a novel mechanism for dysregulation of known driver genes

MM has been reported to have oncogenic dependencies with recurrent mutations affecting 80 known driver genes (**Suppl Table 4**) [10, 19]. We therefore investigated the possible role of rare SVs both in recurrent and rare SV events as a novel mechanism for targeting and dysregulating known myeloma driver genes recurrently involved by single nucleotide variation (SNV). Considering patients without recurrent SNVs in driver genes, we found 48 rare SVs in 45 (6%) patients involving 29 of the 80 known MM drivers. Importantly, twenty-four of the driver genes impacted by rare SV had expression changes in line with their expected role in MM (i.e. duplication associated with overexpression of oncogenes and deletions associated with underexpression of tumor suppressor genes in the 4^th^ and 1^st^ quartiles, respectively), with 15 patients displaying outlier expression in impacted genes (**Fig 3A-B)**. Among these events, we noted gene overexpression mediated by rare focal tandem duplication or templated insertions on *KRAS* (n=1), *NRAS* (n=1) and *PIM1* (n=2), with outlier expression in the patient with *KRAS* impacted and one patient with *PIM1* impacted (**Fig 3B-C; Suppl Table 5**), supporting the potential driver role of these SVs, and providing an alternate mechanism of dysregulating known driver genes. Furthermore, the potential driver nature of these rare events is further supported by the fact that these genes are rarely involved by SVs outside the expected types (e.g., duplication type SVs targeting oncogenes and deletion type SVs targeting tumor suppressor genes; 15/33 expected SV-types with outer-quartile expression vs 5/13 non-congruent SV-types with outer-quartile expression, p < 0.0001, using Fisher Exact). Importantly, in the absence of SNVs, these events would not be detected using whole exome sequencing and/or targeted panel approaches, highlighting the importance of WGS in order to comprehensively capture the genomic heterogeneity of each individual patient.

**Figure 3:**
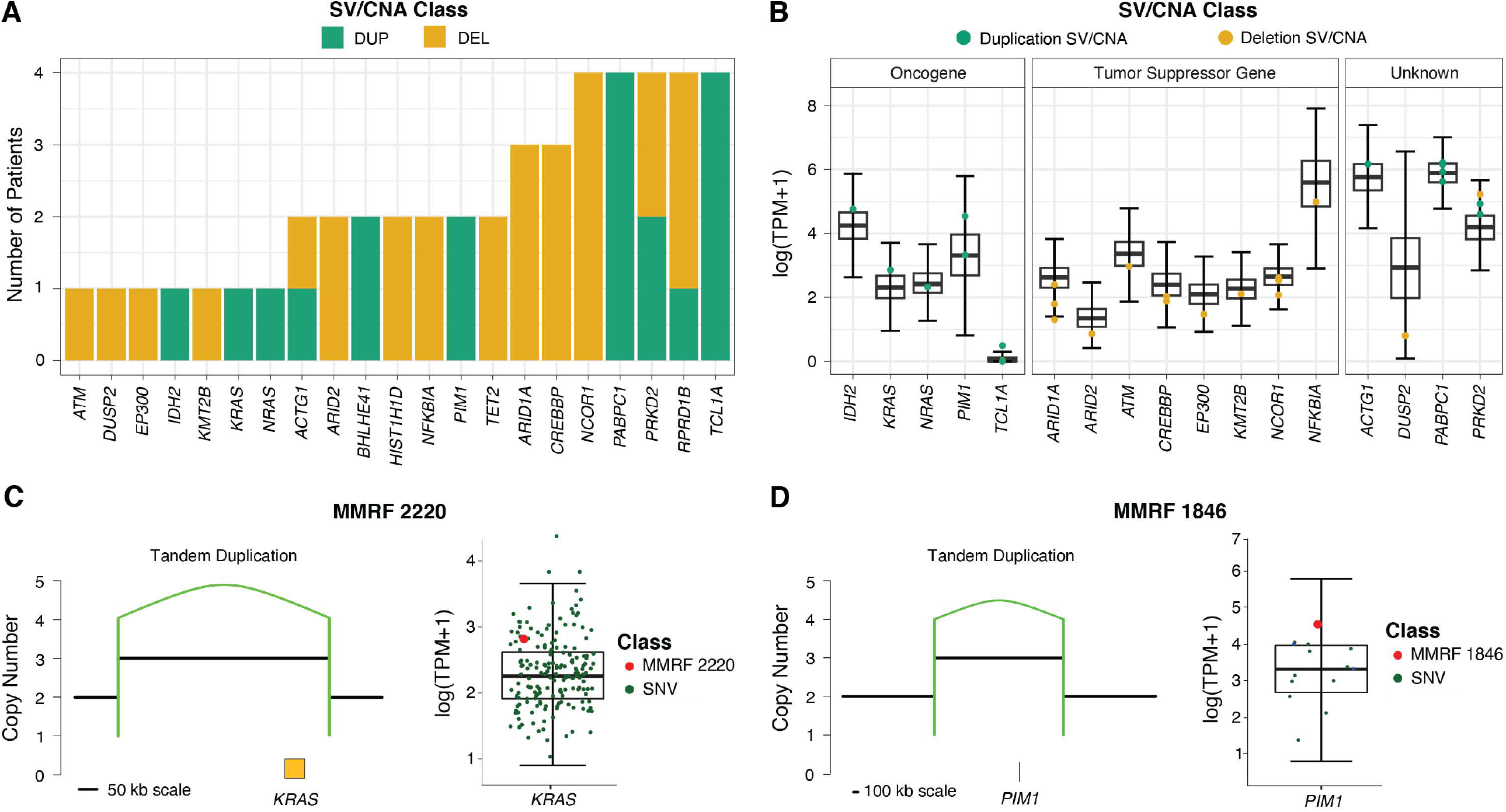
SVs as alternate mechanisms for targeting known multiple myeloma driver genes. **A)** Panel of driver genes classically dysregulated by SNV point mutations focally duplicated of deleted by SVs and CNAs in patient cohort, showing oncogenes affected by duplicated, tumor suppressor genes by deletions, and both duplicated and deletions impacting genes with undetermined impact. **B)** Expression of genes impacted by rare SVs, denoted by colored points and CNA impact. **C-D)** Examples of rare SV targeting known multiple myeloma driver genes. **C)**Tandem duplication SV increasing copy number of *KRAS*, with associated increase in gene expression measured in the 4^th^ quartile (z-score >2). **D)** Tandem duplication SV increasing *PIM1* copy number, with associated increase in gene expression in the 4^th^ quartile (z-score >2). In **C)** and **D)** the minor allele was retained.

### Chromothripsis with and without hypergains

Chromothripsis in MM can present with different genomic and copy number patterns. Overall, we identified two main profiles: 1) multiple deletions with copy number profile oscillating around either monoallelic loss or gain (67.5%, 75/111); 2) focal segments amplified multiple times (32.5%, 36/111). The proportion of rare SV and fully rare events were not different between the 2 chromothripsis patterns (p>0.05, Wilcoxon test). Focusing on the latter, we investigated the frequency of focal (<3 Mb) copy number increases with 3 or more additional copies (i.e. total CN ≥5), which we define as hypergain events. With increasing extra CN, chromothripsis was increasingly responsible for most of the hypergains compared to other SV classes (226/268; 84%) (**Fig 4A**). Focal hypergains associated with chromothripsis were studied in three possible SV patterns: recurrent SVs as part of recurrent chromothripsis events, rare SVs as part of a recurrent chromothripsis event, and rare SVs as part of fully rare chromothripsis events. The proportion of rare and recurrent SVs in recurrent chromothripsis events leading to hypergains was comparable (94/226; 41.6%) and was responsible for the majority of hypergains in rare SVs (94/97; 97%) (**Fig 4B**). Testing the expression of genes involved by rare SVs, within recurrent chromothripsis events, showed clear overexpression of 180 genes, 6 of which do not have a defined driver role in MM including known cancer drivers *SOS1, NTRK1, and FCRL4* (z-score >2; **Fig 4C, Suppl Fig 2, Methods**). Importantly, there is no difference in clinical outcomes between patients with hypergain and non-hypergain chromothripsis, with both groups displaying poorer outcomes compared to patients without chromothripsis (**Fig 4D**).

**Figure 4:**
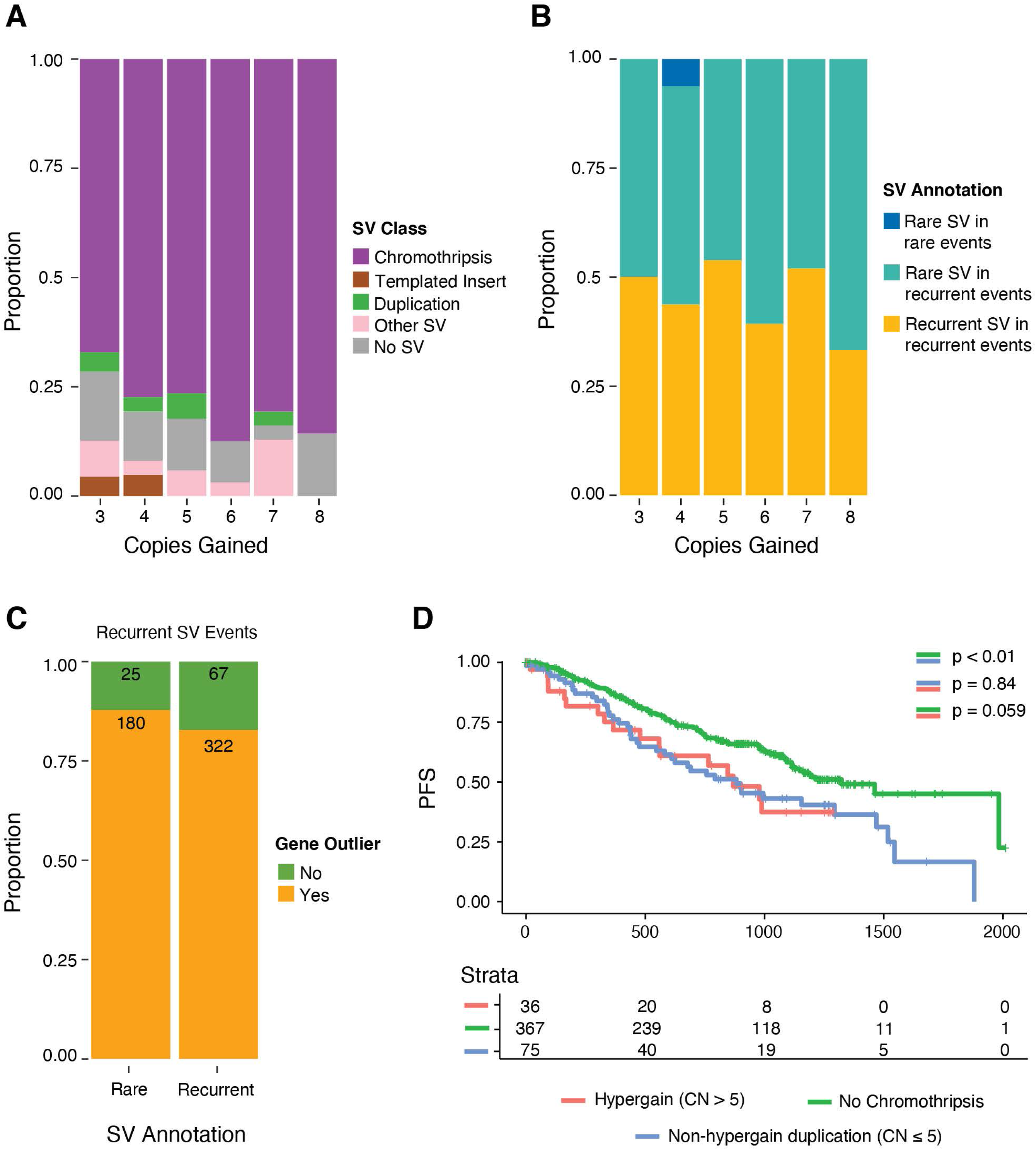
Chromothripsis can cause multiple focal copy number gains associated with increased gene expression. **A)** Chromothripsis dominates as the primary SV class to cause focal copy number increases beyond 3 additional copies – here defined as hypergains. **B)** Rare SVs within recurrent chromothripsis events cause the majority of hypergains. **C)** Of all genes within hyepergain regions affected by rare chromothripsis SVs, the majority exhibit gene expression within the 4^th^ quartile and beyond. **D)** Patients with hypergain chromothripsis display a shorter progression free survival (PFS) compared to patients without chromothripsis. PFS is not different between patients with non-hypergain duplication (CN ≤ 5) and hypergain chromothripsis events.

### Impact of fully rare events and implications on disease biology

To infer the biological impact of each rare SV and distinguish their role between a passenger or a potential driver, we investigated their impact on gene expression both in a direct manner, where an SV occurs within a gene body, and in an indirect manner where they function through transcriptional deregulation via superenhancer hijacking. SVs were considered to involve a gene if either breakpoint intersected the gene region and/or occurred up to 1 Mb away, to take account of distal relationships within a topologically associated domain (TAD) [14, 20]. Distinct duplications and deletions have been reported to recurrently dysregulate the interaction between gene-enhancers by remodeling TADs [4, 5, 21]. To determine SV class-specific breakpoint enrichment in relation to distance from genes, a re-shuffle permutation was performed to create a random background model for each SV class and gene expression direction (i.e., up vs down regulation). Extending on methods developed to test the penetrance of rare germline events, genes paired with rare SV’s were considered affected if the gene expression was outside a gene specific outlier z-score of +/- 2 [22]. Across 591 patients, with available WGS and RNA-seq, we observed a significant association between rare SV events and outlier gene expression. Genes within recurrently affected areas (SV hotspots, GISTIC CNAs, and canonical Ig translocations), were excluded from this analysis, as by definition, a rare SV never involves a recurrently aberrant region.

Across all SV classes and SV to gene outlier distances, we saw a variable distribution of the number of gene outliers involved in each individual SV in fully rare SV events (**Fig 5A**). Notably, rare templated-insertion events affected the most gene outliers per event. Against a background model, rare templated insertions were enriched in overexpressing outliers within the body of genes and up to 1 Mb away (**Fig 5B**). Rare duplication class SVs were enriched in overexpressing outliers within the gene body, in genes 100 kb of the breakpoint and up to 1 Mb away. Rare inversion class SVs were enriched in overexpressing outliers of genes 100kb and 1Mb away. Rare translocations were associated with overexpressing outliers 1 MB away. Rare complex SVs were enriched within the gene body of underexpressing outliers, while deletion SVs were enriched in gene outliers with decreased expression within the gene body, 100 kb away, and up to 1 Mb away (**Fig 5B**). A total of 201 (34%) patients had at least 1 rare SV event associated with gene expression outliers (**Fig 5C**). Additionally, rare SVs in recurrent SV events were enriched within chromothripsis events associated with overexpressed outliers across all gene to SV distances and associated with under-expressing outliers in 100 Kb and 1 Mb relationships (**Suppl Fig 3**). Taking in aggregate all gene expression outliers with SV association up to 1 Mb away, gene ontology pathway analysis (**Methods**) showed that the involved gene outliers are involved in key cell cycle pathways and proliferative Rac mediated signal transduction responsible for ERB2-mediated malignant transformation and metastasis, in addition to involving B-cell development genes such as *PRKDC*, and *POU2F2* (**Fig 5D, Suppl Table 6**) [23, 24].These data suggest that a fraction of rare SV are actively involved in shaping the tumor genomic profile and therefore they are likely not passengers.

**Figure 5:**
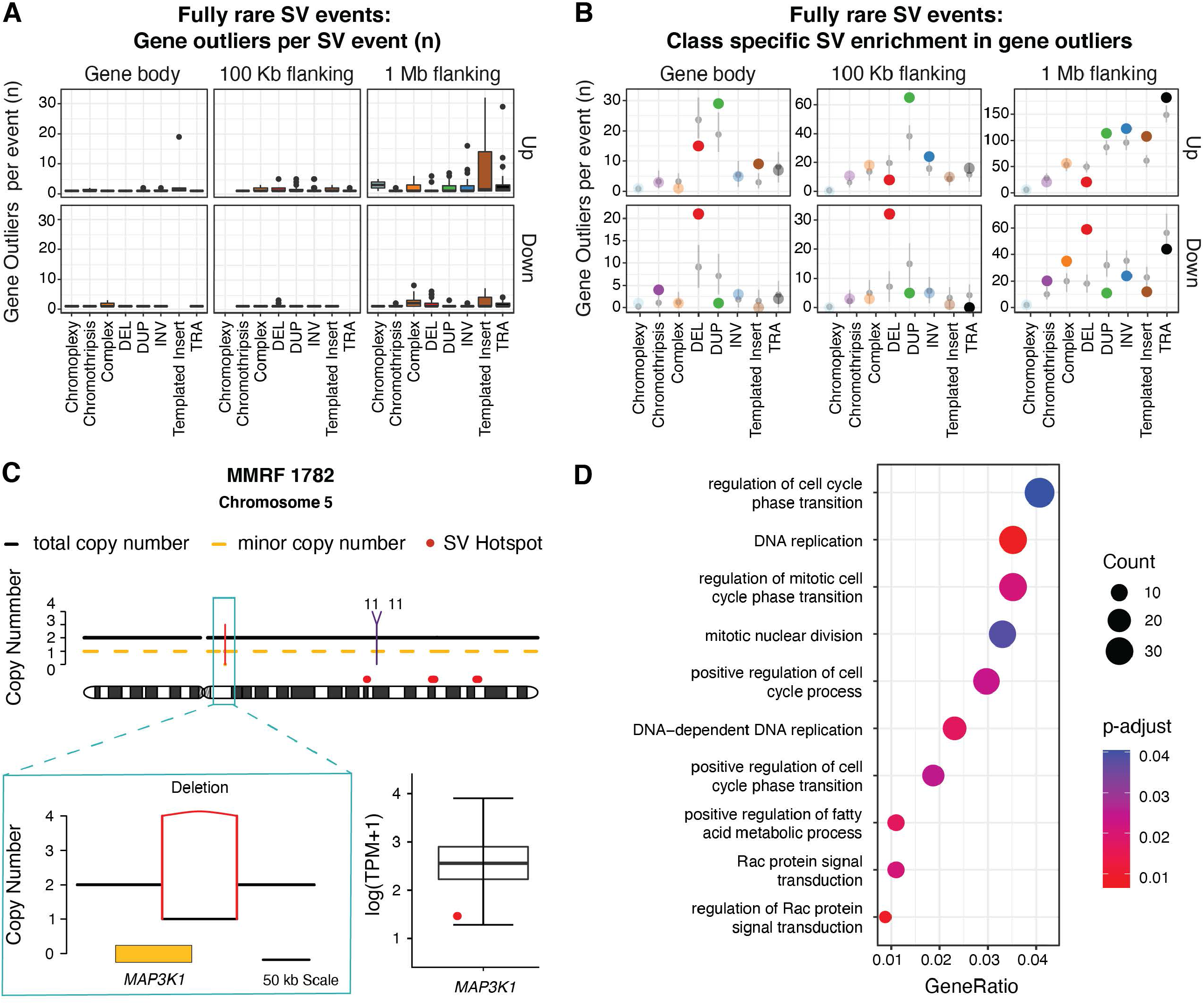
Impact of fully rare SV events on gene expression. **A)** Number of gene outliers (z-score >2 or < -2) in rare SV events subdivided into SV classes and proximity from SV to gene outlier. **B)** Class specific rare SV enrichment in relations gene to SV proximity, subdivided into over-expressing outliers (z-score >2) and under-expressing outlier (z-score < -2). Solid points represent significant enrichment against background premutation rates. Translucent points represent non-significant relationship against background premutation rates. **C)** Example of rare SV event involving a tumor suppressor gene. Single focal rare deletion SV bisecting *MAP3K1* gene with associated low expression in the 4^th^ quartile (z-score < -2), in patient MMRF 1782. **D)** Gene ontology pathway analysis of all gene outliers with relationship to fully rare SV events across all classes and SV to gene outlier distance.

### Rare structural variants and superenhancers

Of the enriched SV to gene outlier relationships, translocation, duplication and templated insertion events were enriched in gene outliers up to 1 Mb distance were of particular interest for their possible involvement in expression regulation through distal regulatory elements. Translocation class SVs have been known to be involved in superenhancer hijacking, leading to up-regulation of distinct oncogenes in MM [25]. This known mechanism is particularly enriched among templated insertions and recurrently affect important oncogenes such as *CCND1, MYC, TENT5C*, and *MCL1* [26]. While these associations have been extensively explored across recurrent SV, the interaction between rare SV and superenhancers has never been investigated. To do so we focused on rare SVs within fully rare SV events, modelling SV breakpoint density to the nearest known MM superenhancer up to 10 Mb [20]. To determine if this association was higher than what would be expected by chance, rare SV breakpoints were re-shuffled, adding a random length between 10-20 Mb to the original position, and proximity to nearest superenhancer was re-calculated to develop a random density distribution.

Overall, rare duplications, translocations and templated insertions were significantly enriched within and/or up to 1 Mb from superenhancer regions, when comparing all other SV classes. Next, we determined if SV breakpoints occurred in a clustered or random manner with regard to superenhancer proximity. To determine this, we first developed a random background model by repositioning templated insertion, translocation, and duplication SV breakpoints 10-20 Mb of the original position in a re-shuffle permutation model (**Methods**). Compared to this randomized background model, only templated insertions emerged as significantly enriched within 1 Mb of superenhancers (p <0.001, Fisher Exact, **Fig 6A**). Considering genes that were directly involved with enriched rare SVs involved SE regions directly and up to 1 Mb distance, and genes within 1 Mb of affected superenhancers, we found 303 gene expression outliers (z-score >2 or <-2). Overall, 26% (127/494) of rare duplications associated with superenhancers were also associated with gene expression outliers, 207 genes being over-expressed (e.g. *IDH2*) and 57 under-expressed, with a median of 3 genes affected per rare duplication event (1-9 gene count range). 36% (65/179) of rare translocation events were associated with outlier genes, with 123 outliers being overexpressed and 33 under-expressed, with a median of 2 genes affected per rare translocation event (1-12 gene count range). 27% (22/82) of rare templated insertion events associated with superenhancers were associated with outlier genes, with 162 being overexpressed (e.g. *IRF6*) and 13 under-expressed, with a median of 21 genes affected per templated insertion event (1-30 gene count range) (**Fig 6B**). The association between rare single and complex SV with superenhancers and gene over expression support the idea that a fraction of these events are not passengers but rather play a potential driver role.

**Figure 6:**
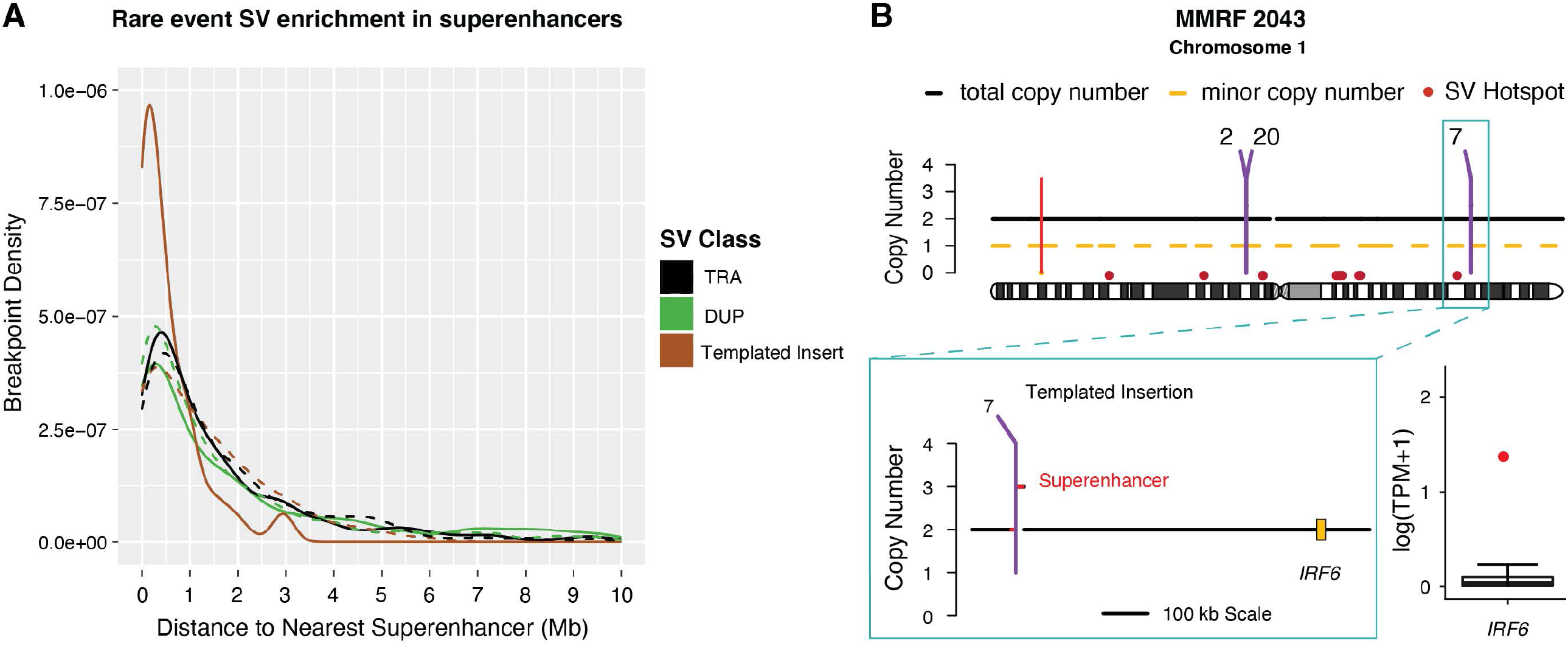
Rare SVs involving myeloma superenhancers. **A)** Fully rare templated insertion class SV events were enriched within 1 Mb of known myeloma superenhancer regions (SE). Solid line depicts observed SV breakpoint density. Dashed line represents breakpoint density expected by permutations background rates. **B)** Example of rare templated insertion SV bisecting and duplicated an SE region. Within 1 Mb downstream of the affected superenhancer, *IRF6* displayed high outlier expression in the 4^th^ quartile (z-score >2).

## DISCUSSION

In recent years, DNA sequencing and expression studies have allowed the description of several new myeloma genomic and clinical subgroups. However, clinically, patients within groups harboring similar or even identical recurrent genomic features present with significant clinical heterogeneity. This clinical reality suggests that our understanding of the MM biology is incomplete, and this might be partially due to unknown genomic drivers contributing to significant disease heterogeneity in each individual patient. WGS has allowed us to expand our catalogue of recurrent genomic drivers, in particular, the characterization of CNA and SVs not involving Ig loci. Nevertheless, several somatic SV events remain unexplained and not associated with any recurrent gene or biological mechanisms known to play a driver role in MM pathogenesis. In this study, we have analyzed a large series of NDMM to decipher the role of rare SV events in NDMM in particular by their impact on gene expression. We focused on rare SVs specifically for two main reasons: 1) the frequency of rare SV events across the genome is low but they have the potential for significant biological impact compared to other genomic features such as SNV and indels, where the great majority are neutral and phenotypically inconsequential, and 2) Unlike SNV on driver genes, somatic SV and CNA have rarely been found in normal cells across different tissues, suggesting that SV and CNA are essential of gene deregulation, selection and tumor progression [22, 27-29]. These two findings suggest that the fraction of passenger SV is likely to be lower than what is seen with SNV and indels. Based on this hypothesis, we applied an analytical workflow that is similar to what has been previously performed to define the impact of rare genetic events [22]. Three rare SV categories were identified. The first group was characterized by fully rare SV events involving genes without a clear and known driver role in MM. Across the entire cohort, 264 patients (35%) had at least one fully rare SV linked to gene expression outliers. In addition, we found a significant enrichment of duplication, translocation and templated insertions at known MM superenhancer regions supporting the link between rare SV and gene expression outliers. The second group is composed of genes without any known driver role in either MM or cancer generally that were affected by multiple focal amplifications. The majority of these events were caused by chromothripsis (226/268; 84%) with 87% (180/205) of genes overexpressed as a consequence of multiple gains associated with rare SVs in recurrent chromothripsis events. This finding demonstrates that rare genes can be involved by distinct and complex events actively impacting individual tumor gene expression. The third group comprised of rare SVs involving known MM driver genes that have been previously shown to be involved by mutations, but not CNA or SV. Interestingly, 45 patients had at least one of these events, and in 33% (15/45) of patients these events impacted gene expression resulting in expression within the 1^st^ or 4^th^ quartile. This is an important finding, as some of these genes are potentially targetable and therefore expand the number of patients who might benefit from future targeted therapies. Taken together these data suggest that genomic and transcriptomic profiles in a significant fraction of patients are affected by rare SV events.

While larger cohorts of WGS are needed to identify new hotspot and driver genes significantly enriched for SV, the picture that emerges from this data is that the heterogeneity between individual patients is also driven by rare SV events, not easily identifiable using standardized statistical approaches based on prevalence and frequency. Furthermore, some of the observed rare SV events may in fact be deemed recurrent in larger datasets with better statistical power. The definition and characterization of these events is not only important for deciphering the individual patient heterogeneity, but might have significant implications for the development of individualized treatment strategies and in our understanding of mechanisms of resistance to novel targeted therapies.

## METHODS

### Patient Cohort and Sample Processing

WGS and RNA-sequencing of bone marrow biopsies were utilized from NDMM patients enrolled in the CoMMpass study (NCT01454297; IA13). Processing of sequencing data was conducted as previously described in the preceding publication [9]. In brief, WGS samples underwent low-coverage long-insert sequencing (median 4-8X). Paired-end reads were aligned to human reference genome (USCS hg19) with the Burrow Wheeler Aligner (BWA; v0.7.8). CNA were identified utilizing tCoNuT (https://github.com/tgen/MMRF_CoMMpass/tree/master/tCoNut_COMMPASS) and externally validated with controlFREEC as previously described [9]. SV identification was performed by two SV callers, DELLY(v0.7.6) and Manta(v1.5.0). The final SV catalogue was obtained by quality filtering calls and validating by copy number alterations as previously described [9]. Annotation of SVs as being involved in complex SV events or as stand-alone SV breakpoint pairs was conducted. SV considered to be causative of a copy number alteration were logged when the SV breakpoint occurs within 50 Kb of a copy number variable start or end position.

RNA-seq data with a target coverage depth of 100 million reads were utilized from 591 patients. Paired-end reads were aligned to the human reference genome (UCSC hg19) utilizing the STAR aligner (STAR; v2.3.1z). Transcripts were normalized to TPM utilizing Salmon (v7.2).

### Defining Rare Structural Variants

Identification of recurrent SVs occurred in a step-wise fashion following an SV involvement hierarchy: 1) SVs involved in canonical myeloma translocations occurring between Ig loci (*IGH, IGK*, and *IGL*) or their partner genes (*CCND1, CCND2, CCND3, MAF, MAFA, MAFB*, and *NSD2*), 2) SVs occurring within or 100 kb from SV hotspot boundaries, and 3) SVs responsible for CNA in the patient cohort, as identified by the GISTIC algorithm. SV hotspot and recurrent CNA region catalogues are reported with our previous publication[12]. We considered the varying nature of recurrent CNA regions involving GISTIC peaks with SV breakpoints as well. CNA regions involving GISTIC peaks vary in size. Both focal (<3Mb) and large CNAs overlapping GISTIC peaks whose breakpoints were within 50kb of an SV were considered causal, involved, and recurrent. Any SV breakpoint involved in at least one of the above event categories defined the SV event as a recurrent event. Therefore, any remaining SV events were defined rare, not involving any known genomic region clinically defined in myeloma disease progression or targeted at significant rates across the patient cohort under investigation.

### Gene Expression Effects of Rare Structural Variants

Genes within SV deletion and duplication class were considered affected by the SV by default. Additionally, genes overlapping part of a duplication and deletion class SV, as well as those occurring up to 1 Mb away were considered in analysis. Genes up to 1 Mb away from inversions and translocations without copy number support were also considered. All SV events, single and complex, were removed from analysis if at least one breakpoint in the event occurred within 1 Mb of any Ig loci, to remove strong immunoglobulin associated enhancer effects. Genes involved in recurrent events were removed from consideration in rare SV analysis. Each rare SV breakpoint was considered individually, cataloguing breakpoints occurring within the gene body, 10-100 Kb, 100 Kb -1 Mb of genes with outlier expression. To examine the relationship between rare SV with outlier expression was analyzed. Extending on methods developed to test the penetrance of rare germline events, genes paired to rare SVs were considered affected if the gene expression was above a gene specific outlier z-score of +/- 2 [22]. Gene outliers were defined as patient specific gene expression with a z-score of +/- 2. To determine if there is a class specific interaction with over-expressed and under-expressed gene expression outliers, a re-shuffle permutation was run on observed counts to model a random distribution of SV breakpoints. If the observed rates of class-specific SV to gene proximity was above the 97.5% CI defined by the random model, the SV class was defined as enriched, having a relationship to gene expression outliers. Cancer related gene data sets were pulled from Memorial Sloan Kettering’s FDA recognized tumor mutation database OncoKB and COSMIC’s Cancer Gene Census, two publicly available repositories [30, 31]. Myeloma driver gene list was pooled from previous publications on driver gene discovery in multiple myeloma [10, 19].

### Superenhancers Involved by Rare Structural Variants

To investigate the relationship and possibility of rare SV influencing gene expression in an indirect manner, we analyzed the relationship between rare structural variants and superenhancers [20, 32, 33]. We first observed SV breakpoint density, up to 10 Mb from the start and end of each superenhancer, considering the closest superenhancer to each breakpoint of the SV. For complex events where multiple breakpoints affect the same superenhancer, only 1 breakpoint per patient was counted. Because we are interested in SV event-to-enhancer relationships, not the number of breakpoints within each superenhancer, the analysis was performed this way to prevent artificially inflating SV involvement in superenhancers. To determine from which distance rare SVs target superenhancers, a re-shuffle permutation was performed randomly shifting the original breakpoint +/- 10-20 Mb, to model a random distribution of SV breakpoints. To determine SV classes that were enriched within 1 Mb of superenhancers, a Fisher Exact test was performed on observed vs modeled counts of class specific SVs within 1 Mb of superenhancers.

## ACKNOWLEDGMENTS

This work was supported by the Paula and Rodger Riney Multiple Myeloma Research Program Fund, the Multiple Myeloma Research Foundation (MMRF), the Perelman Family Foundation, a Memorial Sloan Kettering Cancer Center National Cancer Institute (NCI) Core Grant (P30 CA 008748), and by a Sylvester Comprehensive Cancer Center NCI Core Grant (P30 CA 240139). FM is supported by the American Society of Hematology (ASH).

GJM received grant support through a Translational Research Program award from the Leukemia & Lymphoma Society (6020-20).

## DATA AND SOFTWARE AVAILABILITY

All the raw data used in the study are already publicly available (dbGap: phs000748.v1.p1). Analysis was carried out in R version 3.6.1. The full analytic workflow is reported in **Supplementary Data 1**. All other software tools used are publicly available.

## Notes

**CONFLICT OF INTEREST STATEMENT** OL has received research funding from: National Institutes of Health (NIH), National Cancer Institute (NCI), U.S. Food and Drug Administration (FDA), Multiple Myeloma Research Foundation (MMRF), International Myeloma Foundation (IMF), Leukemia and Lymphoma Society (LLS), the Paula and Rodger Riney Myeloma Foundation, Perelman Family Foundation, Rising Tide Foundation, Amgen, Celgene, Janssen, Takeda, Glenmark, Seattle Genetics, Karyopharm; Honoraria/ad boards: Adaptive, Amgen, Binding Site, BMS, Celgene, Cellectis, Glenmark, Janssen, Juno, Pfizer; and serves on Independent Data Monitoring Committees (IDMCs) for clinical trials lead by Takeda, Merck, Janssen, Theradex.

All other authors have no conflicts of interest to declare.

### Competing Interest Statement

OL has received research funding from: National Institutes of Health (NIH), National Cancer Institute (NCI), U.S. Food and Drug Administration (FDA), Multiple Myeloma Research Foundation (MMRF), International Myeloma Foundation (IMF), Leukemia and Lymphoma Society (LLS), the Paula and Rodger Riney Myeloma Foundation, Perelman Family Foundation, Rising Tide Foundation, Amgen, Celgene, Janssen, Takeda, Glenmark, Seattle Genetics, Karyopharm; Honoraria/ad boards: Adaptive, Amgen, Binding Site, BMS, Celgene, Cellectis, Glenmark, Janssen, Juno, Pfizer; and serves on Independent Data Monitoring Committees (IDMCs) for clinical trials lead by Takeda, Merck, Janssen, Theradex.
GJM has received funding from National Institutes of Health (NIH), National Cancer Institute (NCI), Multiple Myeloma Research Foundation (MMRF), Leukemia and Lymphoma Society (LLS), Perelman Family Foundation, Amgen, Celgene, Janssen, Takeda; Honoraria/ad boards: Adaptive, Amgen, BMS, Celgene, Janssen; and serves on Independent Data Monitoring Committees (IDMCs) for clinical trials lead by Takeda, Karyopharm and Sanofi.
All other authors have no conflicts of interest to declare.

