## Supplementary Figures for "Impact of rare structural variant events in newly diagnosed multiple myeloma"

**Supplementary Figure 1: Rare structural variant landscape in newly diagnosed multiple myeloma. A)** Counts of rare and recurrent events per patients, across full patient cohort, divided into the main cytogenetic groups. **B)** Count and proportion of rare SVs across all SV events, in all patients, divided by cytogenetic groups. **C)** Counts of rare and recurrent individual SVs within rare and recurrent events. **D)** Proportion of rare SVs per event, reported by complex SV class. **E)** Proportion of SVs annotated based on hierarchical annotation (1. Canonical Tra, 2. SV hotspot, 3. GISTIC CNA, 4. Rare SV) within each complex event. Each column denotes one complex event. The color gradient denotes proportion of SVs annotated per hierarchical tier. Summing proportions within vertical bar will result in a proportion of 1 – encompassing all SVs within event. Horizontal groupings denote event classification.

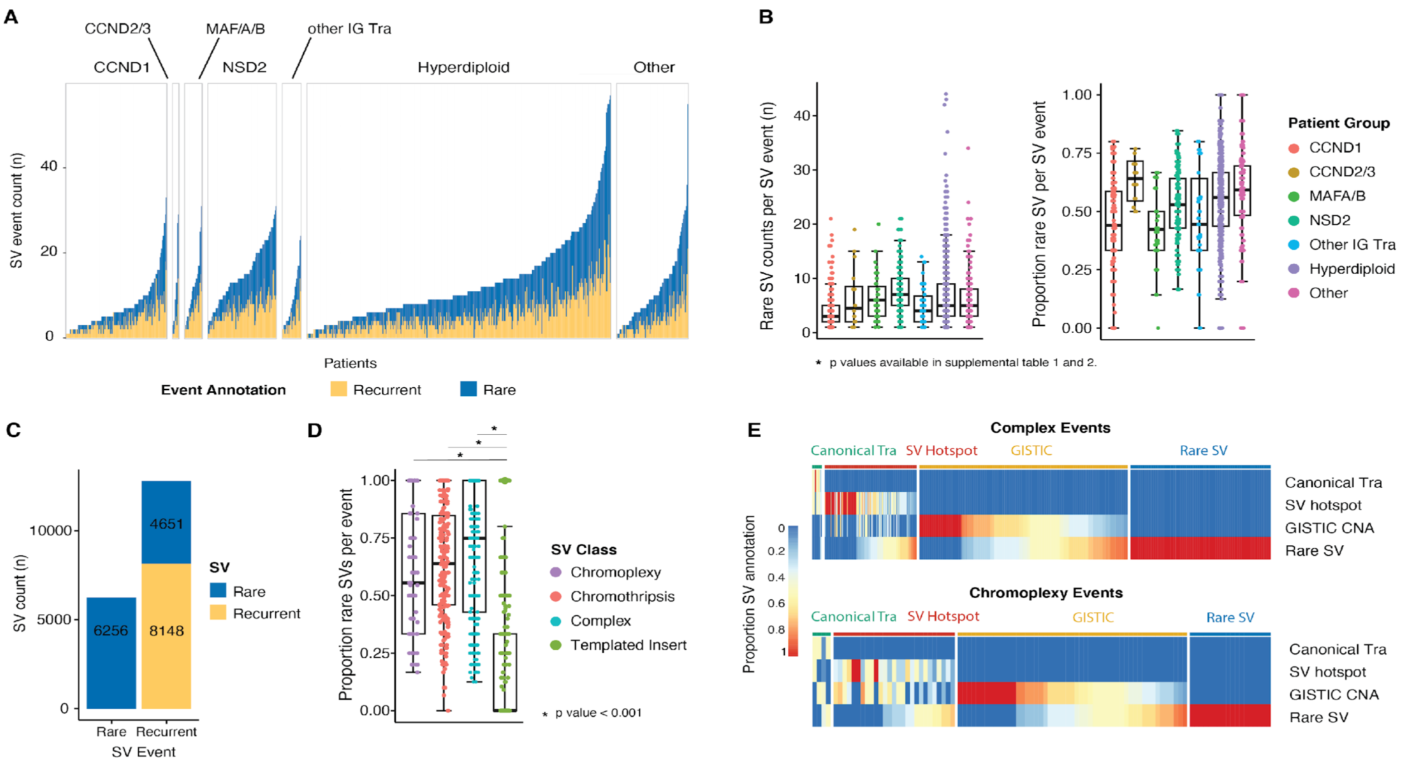

**Supplementary Figure 2: Genes implicated in chromothripsis with hypergains with outlier gene expression.** Green points denote gene directly involved by chromothripsis hypergains with an increase in copy number of 3 or more (total CN ≥5).

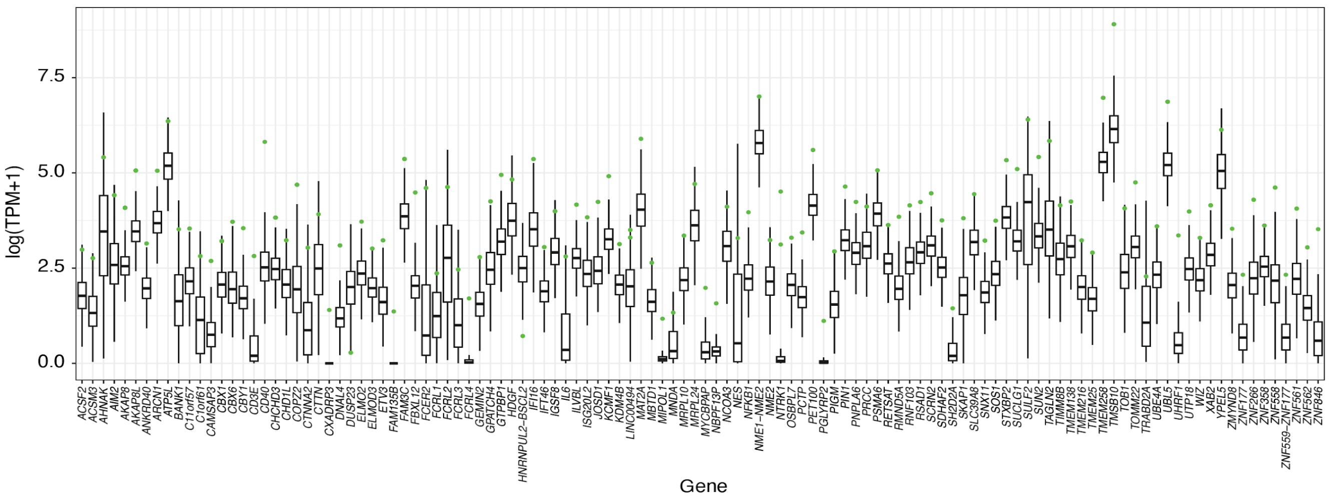

**Supplementary Figure 3: Impact of rare SVs within recurrent SV events on gene expression. A)** Number of gene outliers in rare SV events subdivided into SV classes and proximity from SV to gene outlier. **B)** Class specific rare SV enrichment in relations gene to SV proximity, subdivided into over-expressing outliers (z-score >2) and under-expressing outlier (z-score < -2). Solid points represent significant enrichment against background premutation rates. Translucent points represent non-significant relationship against background premutation rates.

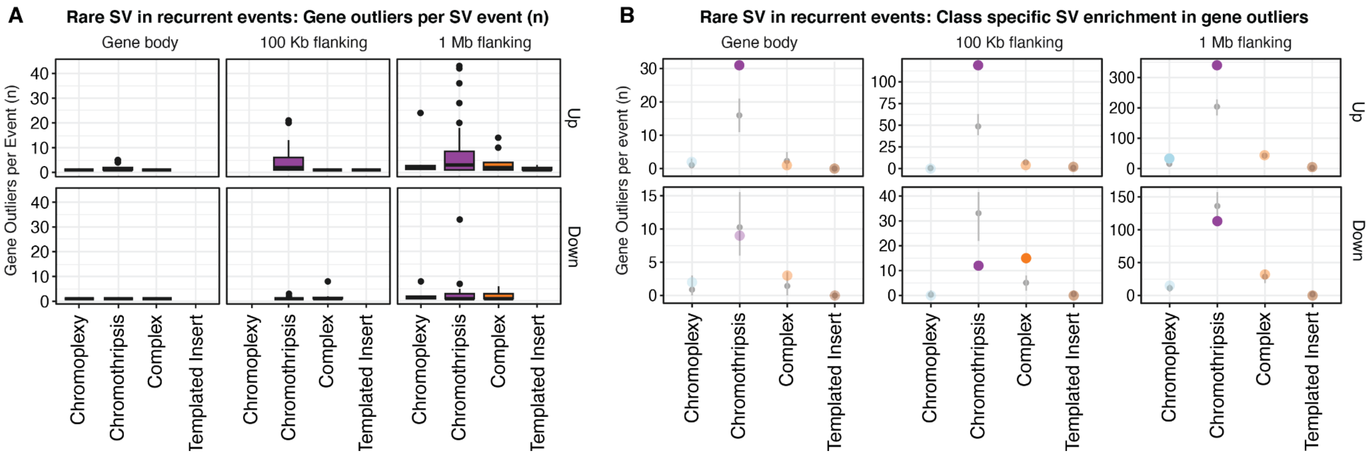
